# Creating Optimal Conditions for OPA1 Isoforms by Western Blot in Skeletal Muscle Cells and Tissue

**DOI:** 10.1101/2023.05.20.541601

**Authors:** Dominique C. Stephens, Margaret Mungai, Amber Crabtree, Heather K. Beasley, Edgar Garza-Lopez, Kit Neikirk, Serif Bacevac, Larry Vang, Zer Vue, Neng Vue, Andrea G. Marshall, Kyrin Turner, Jianqiang Shao, Sandra Murray, Jennifer A. Gaddy, Celestine Wanjalla, Jamaine Davis, Steven Damo, Antentor O. Hinton

**Affiliations:** Department of Molecular Physiology and Biophysics, Vanderbilt University, Nashville, TN, 37232, USA; Department of Life and Physical Sciences, Fisk University, Nashville, TN, 37232, USA; Department of Internal Medicine, University of Iowa, Iowa City, IA, 52242, USA; Department of Biochemistry and Cancer Biology. Meharry Medical College, Nashville, TN, 37208, USA; Department of Cell Biology, College of Medicine, University of Pittsburgh, Pittsburgh, PA, 15260, USA; Central Microscopy Research Facility, University of Iowa, Iowa City, IA, 52242, USA; Division of Infectious Diseases, Vanderbilt University School of Medicine, Nashville, TN, 37232, USA; Tennessee Valley Healthcare Systems, U.S. Department of Veterans Affairs, Nashville, TN, 37212, USA; Vanderbilt University Medical Center: Department of Medicine, Division of Infectious Disease, Nashville, TN, USA

**Keywords:** Muscle Tissue, Mitochondria, Optic atrophy-1 (OPA1), Western Blot, isoforms, isolation

## Abstract

OPA1 is a dynamin-related GTPase that modulates various mitochondrial functions and is involved in mitochondrial morphology. There are eight different isoforms of OPA1 in humans and five different isoforms in mice that are expressed as short or long-form isoforms. These isoforms contribute to OPA1’s ability to control mitochondrial functions. However, isolating OPA1 all long and short isoforms through western blot has been a difficult task. To address this issue, we outline an optimized western blot protocol to isolate 5 different isoforms of OPA1 on the basis of different antibodies. This protocol can be used to study changes in mitochondrial structure and function.

**Tweetable Abstract:** Western blot protocol optimization to visualize OPA1 isoforms.

**Highlights:**

- Protocol for isolating OPA1 isoforms in primary skeletal muscle myoblast and myotubes
- Steps for running isolated skeletal muscle cells from muscle tissue on a gel
- How to collect samples in preparation for western blotting
- Detection of OPA1 isoforms

**Graphical Abstract:** 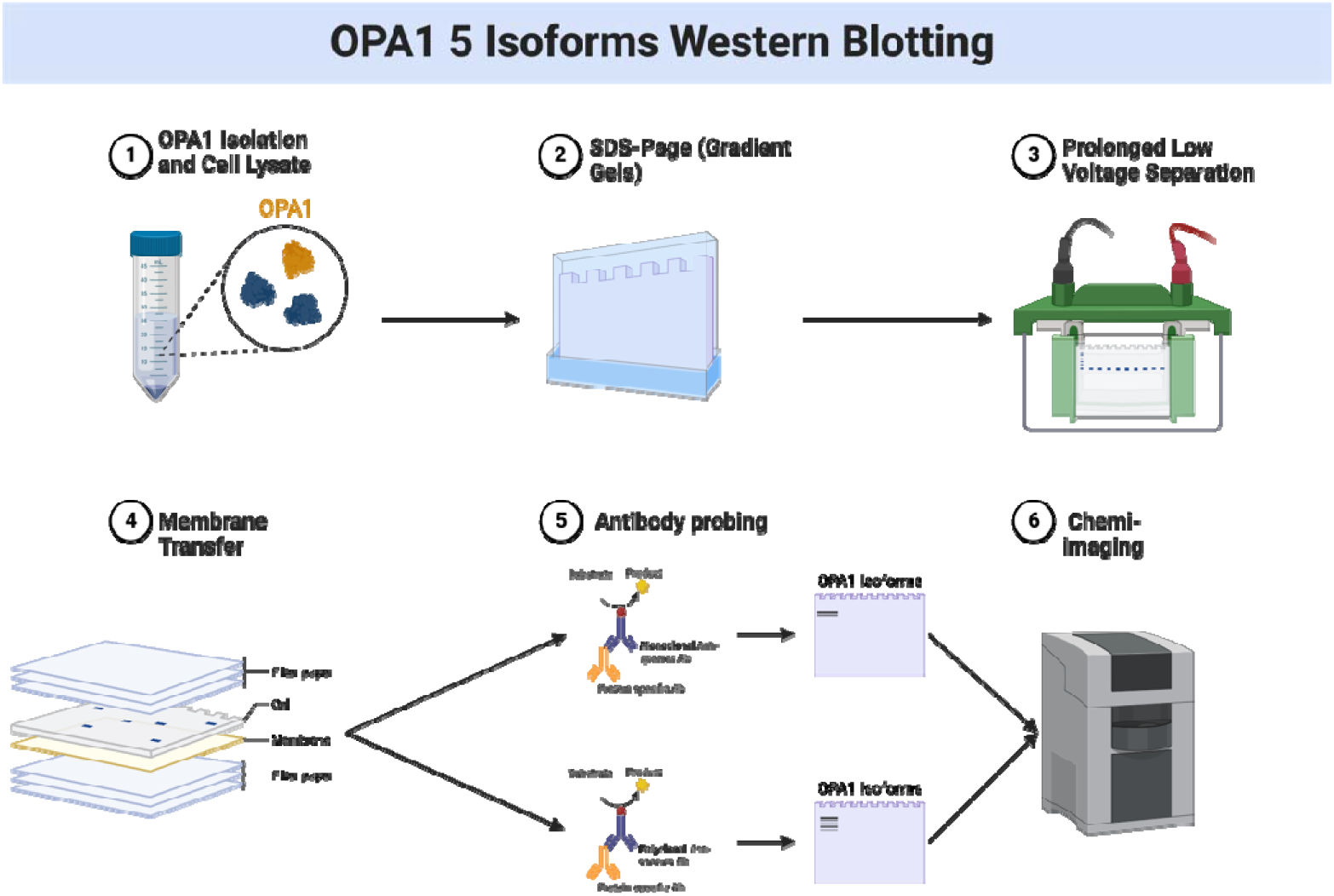

| Key Resources Table |  |  |
| --- | --- | --- |
| Reagent or Resource | Source | Identifier |
| **Antibodies** |  |  |
| Anti-OPA1 antibody (for 2 isoforms) | BD Biosciences | #612606 |
| Rabbit Anti-Mouse IgG H&L | abcam | ab6728 |
| **Experimental Models: Cell Lines** |  |  |
| Primary Myoblast |  |  |
| **Chemicals** |  |  |
| NaCl | Thermo Scientific | 7647-14-5 |
| Tris Base | Millipore Sigma | 77-86-1 |
| SDS | Millipore Sigma | 151-21-3 |
| Triton X-100 | Thermo Scientific | 9002-93-1 |
| Sodium deoxycholate monohydrate | Thermo Scientific | 145224-92-6 |
| EDTA | Millipore Sigma | 60-00-4 |
| Protease inhibitor tablet | Roche | COEDTAF-RO |
| Glycerol | Millipore Sigma | 56-81-5 |
| β-mercaptoethanol | Millipore Sigma | 60-24-2 |
| Bromophenol blue | Thermo Scientific | 115-39-9 |
| DTT | Thermo Scientific | R0861 |
| Glycine | Millipore Sigma | 56-40-6 |
| Methanol | Millipore Sigma | 67-56-1 |
| Tween-20 | Millipore Sigma | 9005-64-5 |
| Dulbecco's Modified Eagle Medium (DMEM) | Thermo Scientific | 11965084 |
| Fetal bovine serum (FBS) | Thermo Scientific | 16000044 |
| PBS | Thermo Scientific | 10010023 |
| Penicillin/streptomycin (Pen/Strep) 100× | Thermo Scientific | 10378016 |
| BSA | Millipore Sigma | 9048-46-8 |
| TBS | Bio-Rad | 1706435 |
| 2X Laemmli sample buffer | Bio-Rad | 1610737 |
| **Quantification and Software** |  |  |
| GraphPad | GraphPad Software, San Diego, California USA | www.graphpad.com |
| Image J | Schindelin et al.5 | https://imagej.net/ |
| **Critical Commercial Assays** |  |  |
| SuperSignal™ West Pico PLUS Chemiluminescent Substrate | Thermo Scientific™ | 34579 |
| **Other** |  |  |
| 100 mm plate | Corning | 353003 |
| Microcentrifuge tube | Thermo Scientific | AM12450 |
| 22 G needle | Amazon | LY-999 |
| Tris-glycine Mini Gels | Novex | XP00100BOX |
| Vertical Electrophoresis System | Bio-Rad | 1658004 |
| Nitrocellulose membrane | Bio-Rad | 1620115 |
| Microcentrifuge | Eppendorf Digital Centrifuge | 5415D |

**Method Summary:** Samples for western blot analysis are isolated from lysed cells, loaded onto a gel, and ran using optimized conditions to better isolate OPA1 isoforms. Samples are transferred to a membrane for incubation and protein detection using OPA1 antibodies.

## Background

Optic atrophy-1 (OPA1) is a GTPase located in the inner membrane of mitochondria and is a central player in mitochondrial fusion ^1^, which is essential for maintaining mitochondrial network morphology and function. OPA1 is also involved in regulating cristae organization, protecting against apoptosis^2^, and maintaining respiratory chain assembly and functionality, mitochondrial membrane potential, and mitochondrial DNA ^1,3^.Furthermore, mutated OPA1 has been linked to a wide range of neurodegenerative diseases, including autosomal dominant optic atrophy ^4,5^. Further research on OPA1 and its functions may lead to new treatments for various diseases.

While humans express eight OPA1 mRNA transcripts by alternate splicing of exons 4, 4b and 5b^6^, mice, mice express 4 variants, isoforms 1, 5, 7 and 8, through alternate splicing of only 4b and 5b ^7^. These isoforms can present as long-form or short-form isoforms associated with different mitochondrial functions across several proteases responsible for processing. In example, in a membrane potential-dependent manner, there is increased fission^8^, long isoforms 1 and 7 are cleaved to short isoforms by OMA1 protease cleaves at site S1 encoded by exon 5 domain and YMLE1L protease cleaves S2 site encoded by exon 5b domain. Isoforms 5 and 8 are processed into only short forms ^7^. Recently, Wang et. al, found a third cleavage site for OPA1, YMEL1 site 3 ^9^. While long-form and short-form isoforms are able to restore cristae structure, mitochondrial DNA (mtDNA) abundance, and energetic efficiency ^6^, long and short isoforms have differential roles. Certain experimental conditions, typically associated with cellular stress such as cold exposure, have been illustrated to cause increased cleavage of OPA1 to short isoforms ^10^. However, short isoforms still play important roles in energetic efficiency, requiring a balance of short and long isoforms ^6^. While traditional monoclonal antibodies allow for seeing these two principal long and short isoforms, they do not allow for additional isoforms to be observed beyond these two ^11^. Recently, researchers found that OPA1-specific isoforms may have functional roles, such as 1 and 7 which have therapeutic potential for diseases caused by mitochondrial dysfunction ^12^. Therefore, it is highly important to be able to efficiently isolate and study OPA1 isoforms. Here, we show optimal conditions for isolating OPA1 isoforms by western blot.

## Materials and Methods

### Buffers

To lyse the cell, we used a RIPA buffer:

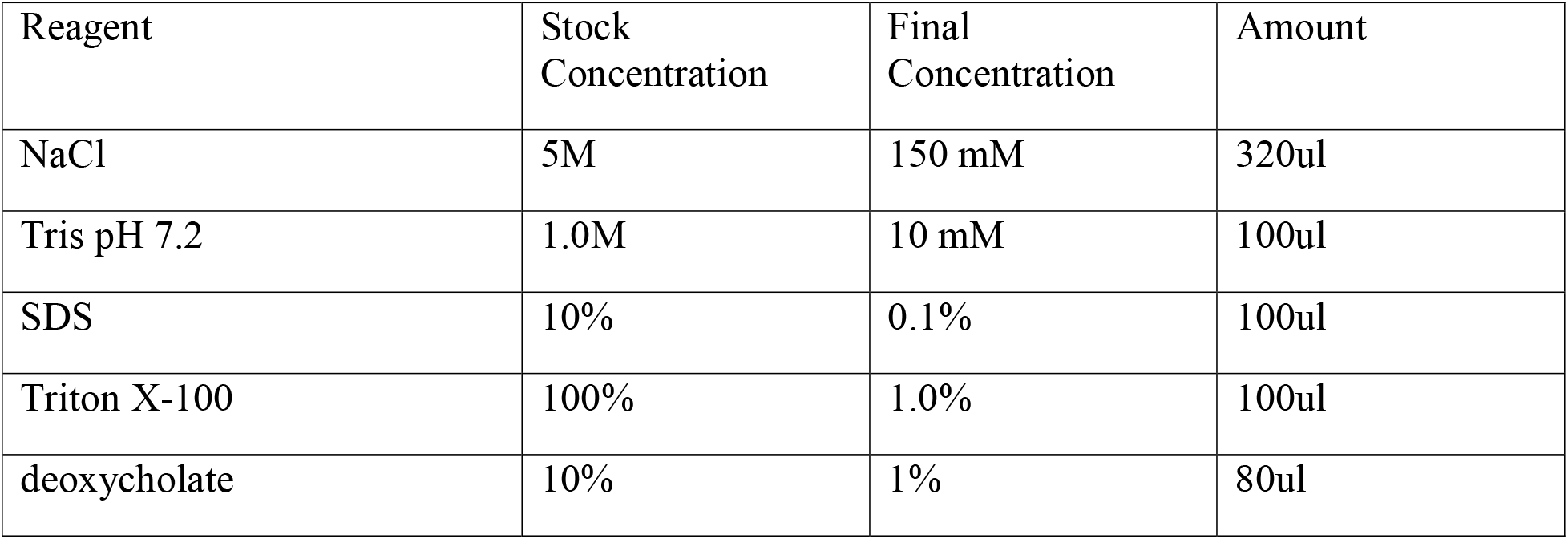
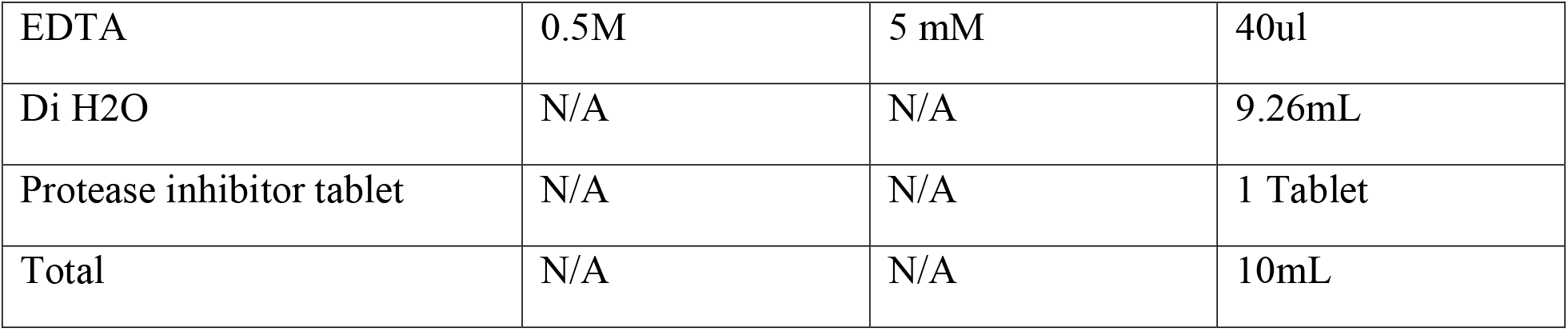

| Reagent | Stock Concentration | Final Concentration | Amount |
| --- | --- | --- | --- |
| NaCl | 5M | 150 mM | 320ul |
| Tris pH 7.2 | 1.0M | 10 mM | 100ul |
| SDS | 10% | 0.1% | 100ul |
| Triton X-100 | 100% | 1.0% | 100ul |
| deoxycholate | 10% | 1% | 80ul |
| EDTA | 0.5M | 5 mM | 40ul |
| Di H2O | N/A | N/A | 9.26mL |
| Protease inhibitor tablet | N/A | N/A | 1 Tablet |
| Total | N/A | N/A | 10mL |

Once cell lysate samples are obtained, we mixed our samples with 6X sample buffer before loading:

| Reagent | Stock Concentration | Final Concentration | Amount |
| --- | --- | --- | --- |
| Tris pH 6.8 | 1.5 M | 0.375 M | 2.5mL |
| Glycerol | 100% | 48% | 4.8mL |
| SDS | N/A | 6% | 0.6g |
| β-mercaptoethanol | 100% | 6% | 0.6mL |
| Bromophenol blue | N/A | 0.06% | 0.006g |
| DTT | N/A | 0.6 M | 0.930g |
| DI H2O | N/A | N/A | 2.1mL |
| Total | N/A | N/A | 10mL |

After the samples were loaded onto an SDS-PAGE gel, we used 1X Tris-glycine buffer as our running buffer:

| Reagent | Stock Concentration | Final Concentration | Amount |
| --- | --- | --- | --- |
| Tris base | N/A | 25 mM | 12.1g |
| Glycine | N/A | 250 mM | 75.06g |
| SDS | N/A | 0.1% | 4g |
| DI H2O | N/A | N/A | 4L |

To transfer to a membrane, we used a 1X Tris-glycine transfer buffer:

| Reagent | Stock | Final Concentration | Amount |
| --- | --- | --- | --- |

|  | Concentration |  |  |
| --- | --- | --- | --- |
| Tris base | N/A | 25 mM | 12.1g |
| Glycine | N/A | 192 mM | 57.6g |
| Methanol | 100% | 20% | 800mL |
| SDS | N/A | 0.02% | 0.8g |
| DI H2O | N/A | N/A | 4L |

The following buffers needed for antibody incubations: Blocking buffer

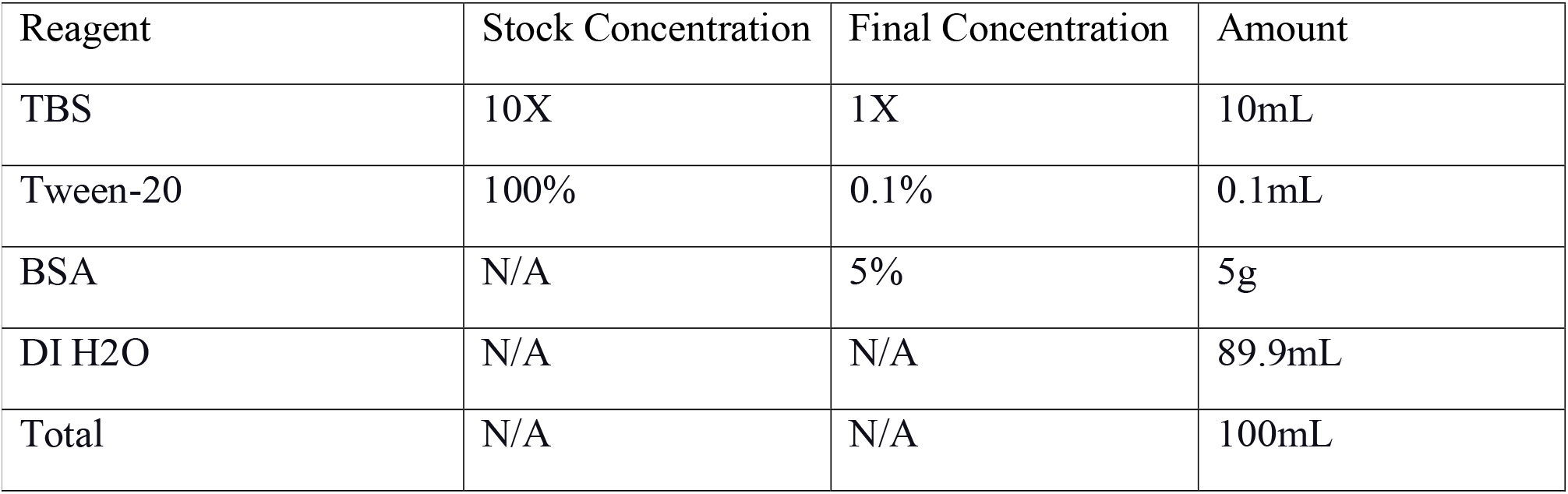

Antibody dilution buffer:

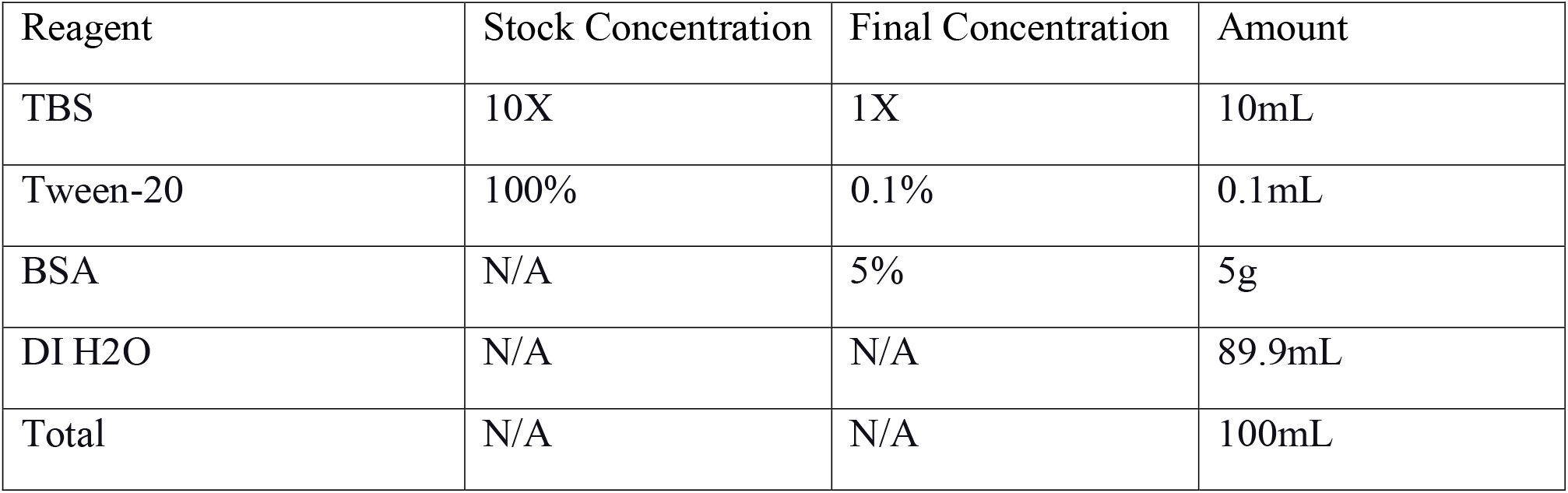

Wash buffer TBS-T:

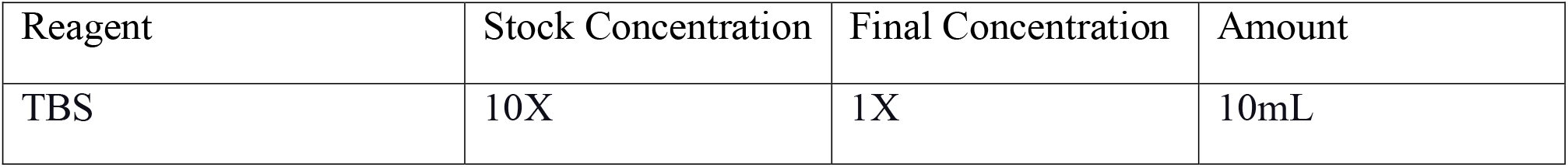
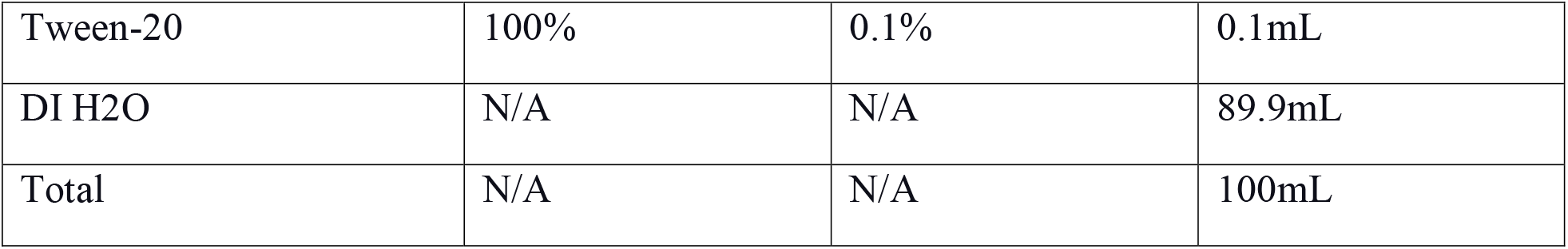

### Western Blot Protocol for Isolating OPA1 Isoforms in Mouse Primary Skeletal Muscle Cells

You can run a two-band blot of OPA1 by running a standard western blot protocol and then blotting for OPA1 (**Figure 1**). However, if you happen to see a change in your OPA1 protein level, we outline a protocol to look at all isoforms of OPA1 in mice (**Figure 2**).

**Figure 1.**
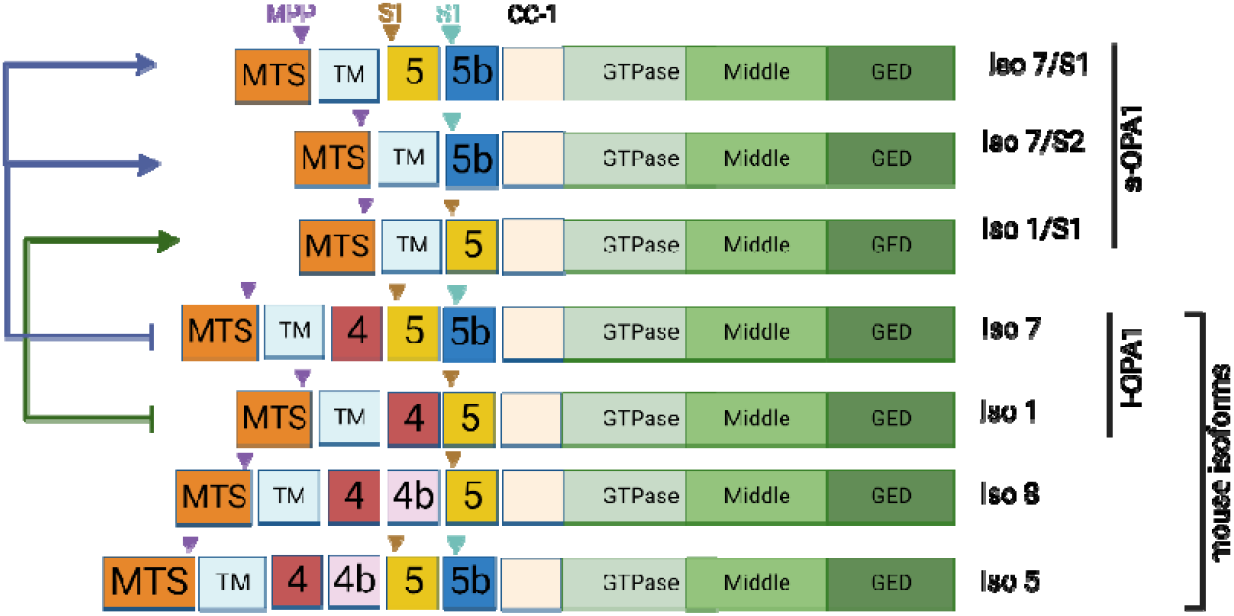
Schematic Representation of the murine OPA1 protein variants. The Opa1 gene has 4 isoforms, isoforms 1, 7, 5, and 8, which are further processed into long (l-OPA1) and short (s-OPA1) forms by S1 and S2 site cleavage by OAM1 and YMEL1 respectively. MIS, mitochondrial import sequence, TM, Transmembrane domain, CC-1, Coiled-coil domain, MPP, mitochondrial processing peptidase, domains 4, 4b, 5, and 5b are encoded by their respective exons 4, 4b, 5 and 5b.

**Figure 2.**
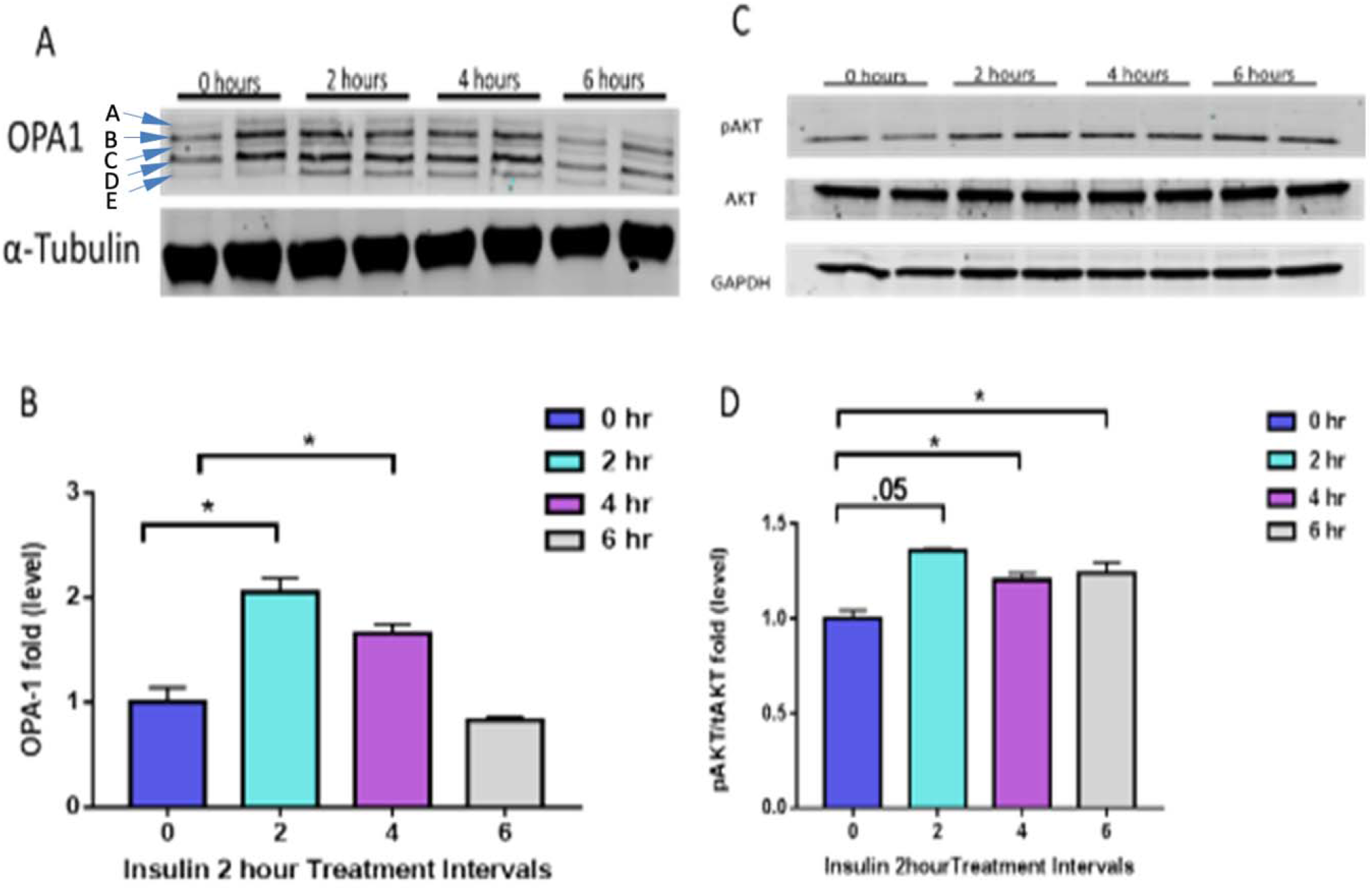
Example of five OPA1 bands with BD Monoclonal anti-OPA1 mouse antibody. Post-2 hours and 4 hours Insulin Stimulation Increases OPA1 levels in C2C12 Myotubes **(A)** Western blot expression of OPA1 and alpha-tubulin, following treatment with insulin for 0 – 6 hours. (**B**) OPA1 levels normalized to alpha-tubulin expression across insulin stimulation intervals. (**C**) Western blot of OPA1, AKT/Protein Kinase B, and phosphorylated AKT, the active form of AKT, following treatment with insulin for 0 – 6 hours. (**D**) AKT and pAKT levels normalized to GAPDH expression across insulin stimulation intervals. N = 6 per treatment with triplicates, and * indicates p-value < .05

1. Day 1:

## CRITICAL

Everything must be chilled in advance

*This section describes how to collect and prepare samples*.

a. Aspirate media
b. Wash cells with ice-cold PBS to remove residual media
c. Aspirate PBS
d. Add RIPA buffer directly to cells.
e. Scrap cells from the plate
f. Transfer scraped cells to a microcentrifuge tube.

**Note:** We used 1 mL of RIPA buffer per 100 mm plate. Scale up or down as necessary.

g. Incubate the microcentrifuge tubes on ice for 10 mins (vortexing every few minutes)

**Note**: Lysates can also be passed through a 22 G needle to aid in solubilization.

h. Centrifuge the microcentrifuge tubes at 7245 RCF for 15 mins.

*This section describes how to run protein samples on an SDS-PAGE gel*

i. Load 20-50 μg of lysate with loading dye (Bio-Rad 2X Laemmli sample buffer and add BME)
j. Run lysate samples on **Novex 4-20% Tris-glycine Mini Gels** (pre-chilled for one hour in cold running buffer before use).

**Note**: You can make your own gels also, but it is important to use a gradient gel, as they provide better separation than normal gel for the close molecular weights of OPA1 isoforms.

**Note**: It is important to ensure that all materials and reagents, especially the Novex Mini Gels, are chilled for an hour prior to use.

i. Run gel for **10 mins at 100 V** (small gel boxes) to get the sample through the wells.

**Note:** For larger gel boxes, 170-200 V can be used, but multiple isoforms may not be seen.

k. Reduce the voltage to **50 V for 5-7 hrs at 4C**.

**Note:** Gels can be run overnight at **35 V at 4C**.

2. Day 2:

*This section describes how to transfer proteins from a gel to a membrane*.

a. Transfer the Tris-glycine gel to a **nitrocellulose membrane** for **12 hours at 35 mV** in a cold room.

### Antibody Incubations

b. Wash the membrane in 1X TBS for 5 mins.
c. Cover the membrane with the blocking buffer.
d. Shake membrane in blocking buffer overnight at 4 L or 1 hr at room temperature.

3. Day 3:

a. Wash the membrane three times with TBST for 5 mins each.
b. Incubate membrane with primary antibody for 1 hr at room temperature,
c. Wash three times with TBST for 5 mins each.
d. Incubate and shake the membrane with the appropriate conjugated secondary antibody for 1 hr at room temperature.
e. Wash the membrane three times with TBST for 5mins each.

### Protein Detection

f. Prepare ECL detection reagent by mixing solutions A & B in a 1:1 ratio.
g. Cover membrane with ECL solution
h. Incubate for 5 mins at room temperature with gentle rocking.
i. Remove the membrane from the solution.
j. Wrap the membrane in plastic wrap, (while eliminating air bubbles)
k. Place the membrane in an exposure cassette.
l. Expose to film for different amounts of time.

### Expected Outcomes

Notably, while OPA1 deficiency causes a loss of mitochondrial morphology it also promotes insulin sensitivity in a fibroblast growth factor 21-dependent manner ^13^. Here we used C2C12 myotubes and, following serum starvation overnight, performed insulin treatment (10 nM/L) in 2-hour increments. To begin with, we performed the above Western blot procedure with the OPA1 BD monoclonal antibody (Figure 2A), which showed a progressive increase of OPA1 levels across 4 hours due to insulin stimulation in C2C12 (Figure 2B). We used 2 different factors to allow for normalization and as an internal control gene, GAPDH and alpha-tubulin. Paralleling this was a similar change in AKT which was additionally consistent across 6 hours of insulin stimulation (Figure 2C-D). Previously in cardiomyocytes, it was elucidated that both 2 isoform bands of OPA1 and 5 isoforms of OPA1 show a similar change in expression due to insulin stimulation ^11^. Here, we show that changes in the total expression of OPA1 can be noted by looking at 5 isoforms of OPA1 as opposed to only long and short isoforms. Given that the AKT pathway aids in regulating Opa1 cleavage ^14^, it was expected that there would be concomitant changes in AKT and the active phosphorylated form of AKT. Furthermore, it is well established that insulin stimulation increases the expression of AKT ^15^. Notably, recent studies in cardiomyocytes have elucidated that insulin stimulation activates the Akt-mTOR-NFκB pathway to increase OPA1 ^11^. We saw a similar increase in AKT as OPA1, marked by significant progressive increases after 2 hours (Figure 2C-D).

We further validated this approach to visualize 5 isoform bands in skeletal muscle. To do this, we compared OPA1 levels following knockdown (smKO/KD) of *Opa1*. In gastrocnemius muscle (Figure 3A) 5 bands of OPA1 were observed, alongside an expected decrease in OPA1 following smKO (Figure 3B). This was similarly observed in soleus tissue (fFigures 4C-D), suggesting this technique is applicable to a range of tissue types and 5 bands of OPA1 reflect protein content in the cell following smKO of *Opa1*.

**Figure 3.**
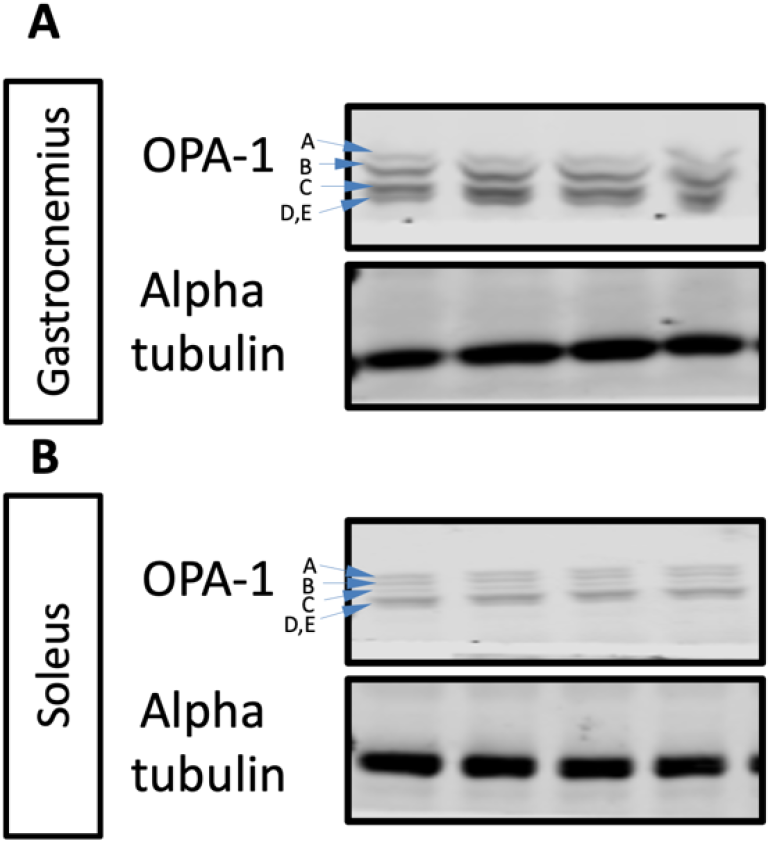
Example of five OPA1 bands with BD Monocolonal anti-OPA1 mouse antibody. **(A)** Western blot expression of OPA1 and alpha-tubulin in gastrocnemius. **(B)** Western blot expression of OPA1 and alpha-tubulin in soleus muscle.

### Quantification and statistical analysis

Bands from the blot are quantified using ImageJ. Statistical analyses were performed using One-way Anova on GraphPad software, significant difference was accepted when P < 0.05.

### Limitations

The main limitation of western blotting is that researchers are confined to using commercially available primary antibodies for detection. Beyond this, cross-reactivity with other proteins or nonspecific binding can lead to false-positive or false-negative results. Here, specific antibodies are utilized depending on the OPA1 isoform desired, but polyclonal antibodies in certain cell types such as cardiomyocytes may not detect modifications in the levels of the OPA1 protein at an early stage ^11^. Beyond that, it is unclear if the number of OPA1 isoforms is dependent on the cell type and if certain models have more or less isoforms. We have only noted 5 isoforms using this protocol, but it may be possible to optimize it to look at all 6 of the known OPA1 isoforms ^6,12^. Specifically, future advances may try to look at relative abundance of specific bands, as the top two bands are hypothesized to be long forms of OPA1 while the bottom 3 bands are thought to be short isoforms ^16^. Elucidating expression of these specific isoforms may be important given their differential role ^6^.

Looking at other proteins such as AKT to confirm associated changes may be important. Finally, it is possible that, depending on the experimental conditions, cleavage of OPA1 to C-terminal fragments is happening causing bands not to occur ^17^. Given that this C-cleavage is dependent on Mfn2, and not common cleavage factors like the metalloprotease OMA1, it can be more difficult to look at upstream factors in such scenarios ^8^.

### Troubleshooting

#### Problem 1

You do not detect any bands on your membrane.

### Potential Solution

This can be fixed through several ways to increase the retention time. One may Increase the primary antibody inoculation time to O/N rocking in a cold room. Alternatively, increase the secondary antibody to no longer than 2hrs. Increase the amount of film exposure time. This lack of detection could be a result of the antibody not having enough time to bind to the proteins on the membrane. Also, we recommended being sure that the primary antibody has not expired, as that would decrease its effectiveness.

#### Problem 2

Your protein did not transfer from the gel to the membrane.

### Potential Solution

This can be checked by using a bromo blue stain on the gel. If bands show up then the proteins were not transferred. Additionally, it is important to ensure that the stacking order is correct to avoid potential issues. Finally, it is important to optimize transfer to avoid weak or undetectable signals, without overtransfer, which can lead to nonspecific binding.

#### Problem 3

You are unable to tell how many isoforms are on the band

### Potential Solution

Smaller separation means a smaller distribution of the different isoforms. We have found that 5-7 hour separation at 50V is key to allowing for all isoforms to be distinguishable. However, tweaking of experimental design to increase retention time may be necessary.

## Resource Availability

### Lead contact

Further information and requests for resources and reagents should be directed to and will be fulfilled by the lead contact, Antentor Hinton.

### Materials availability

All generated materials, if applicable, are created in methods highlighted in the text above.

### Data and code availability

Full data utilized and requests for data and code availability should be directed to and will be fulfilled by the lead contact, Antentor Hinton.

### Financial & Competing Interests’ Disclosure

#### All authors have no competing interests

This project was funded by the UNCF/Bristol-Myers Squibb E.E. Just Faculty Fund, BWF Career Awards at the Scientific Interface Award, BWF Ad-hoc Award, NIH Small Research Pilot Subaward to 5R25HL106365-12 from the National Institutes of Health PRIDE Program, DK020593, Vanderbilt Diabetes and Research Training Center for DRTC Alzheimer’s Disease Pilot & Feasibility Program. CZI Science Diversity Leadership grant number 2022-253529 from the Chan Zuckerberg Initiative DAF, an advised fund of Silicon Valley Community Foundation (to A.H.J.). NSF EES2112556, NSF EES1817282, NSF MCB1955975, and CZI Science Diversity Leadership grant number 2022-253614 from the Chan Zuckerberg Initiative DAF, an advised fund of Silicon Valley Community Foundation (to S.D.) and National Institutes of Health grant HD090061 and the Department of Veterans Affairs Office of Research award I01 BX005352 (to J.G.). Additional support was provided by the Vanderbilt Institute for Clinical and Translational Research program supported by the National Center for Research Resources, Grant UL1 RR024975–01, and the National Center for Advancing Translational Sciences, Grant 2 UL1 TR000445–06 and the Cell Imaging Shared Resource.

